# Molecular characterization of microbes in the center of barnacle footprints (part I): Molecular characterization of microbes in the center of barnacle footprints (part I): a case study for (under)graduate student laboratory research training

**DOI:** 10.1101/2022.11.12.516243

**Authors:** Zhizhou Zhang

## Abstract

There is a barnacle larva settlement model in which barnacle cyprid selectively locates itself only on a specific marine biofilm area that contains specific microbes. That means a local biofilm site with specific characteristics can attract barnacle larva to the maximum. If this is true, a barnacle already growing up shall still press down a chunk of biofilm area where it settles when it was a cyprid. The chunk of biofilm should be at the center of barnacle footprint and may still contain most of the microbes at the attachment site. By this consideration, a group of such chunks of barnacle cement (with about 2mm diameter) was collected from the center of barnacle footprints, followed by genomic DNA extraction, PCR amplification with primers representing prokaryotes, eukaryotes, archaea and fungus, DNA sequencing and species determination. The most abundant 13 species were preliminarily determined (mainly fungi). Whether they are really wanted target microbes largely depends on future investigations on whether they possess some common features that can attract barnacle cyprids. This investigation provided a case study for undergraduate or graduate student to go through a laboratory research process: planning, design, sampling, data collection, experimental techniques, data statistics, and manuscript/report writing.

## Introduction

Mechanisms of barnacle attachment on different marine stratums have been undertaken for over half a century, and vast knowledge on this issue has been gathered. Unfortunately, there is still a distance from a clear understanding on how to prevent its attachment in an environmental friendly way. Knowledge of dynamic interaction between barnacle crypid and microbes plus small creatures in the biofilm is still vague. The relationship between barnacle attachment and local biofilm characteristics is almost still blank till now.

Literature indicates that barnacle attachment and growth contain complex information. The protein sequences in the barnacle cement are unique, different from any known mucoproteins. Though barnacle cyprid has its own preferred sites to attach, those positions that can stably adhere to barnacle cements on the biofilm surface or artificial coating surface can be perpetual sites for barnacle to live.

A batch of studies have been performed in the past many years to analyze the process of barnacle larva settlement. Stanley et al [1] found that crypid settlement of *B. amphitrite* and *B. trigonus* was promoted by biofilms developed at high temperatures (23 and 30°C), but was unaffected (*B. amphitrite*) or inhibited (*B. trigonus*) by biofilms developed at 16°C. This study suggested that biofilm composition or membrane property was selected by barnacle crypid. Lee et al [2] observed that cyprids of Balanus Amphitrite preferred to settle on the intertidal rather than subtidal biofilms, independent of the biofilm bacterial density or biomass, strongly suggesting that barnacle larva settlement may be related with specific bacterial groups in the biofilm. Ahlem et al [3] found that several compounds isolated from the Mediterranean brown alga *Taonia atomaria* have differential antifouling capacity for several bacteria and can inhibit settlement for two barnacle species (*Amphibalanus amphitrite* and *Balanus perforatus*). One study [4] found that periphytic diatoms such as *N. ramosissima* play an important role in larval settlement of the barnacle *A. amphitrite*. The cue in *N. ramosissima* was an Lentil Agglutinin (LCA)-binding sugar chain(s) compound that may be functionally similar to the settlement inducing protein complex (SIPC) [5] from adult shell of the barnacle. Another study [6] reported that the surface-associated cues mediate the settlement and metamorphosis, whereas the waterborne cues could be more significant in locating the destination of larva settlement.

In this study, the question (the relationship between biofilm composition and barnacle larva settlement) was tackled in a new angle. The author guessed that, if the barnacle crypid chooses a specific local biofilm area (with the targeted microbes) to settle down, the crypid either directly tear the biofilm or ‘sit-on’ the local area, then secret the sticky cement around the settlement site. So the targeted microbes have a chance to be pressed beneath the barnacle body for the rest of barnacle life. If this is true, the author will have a chance to pick up those targeted microbes from the barnacle footprints.

## Materials and methods

The iron-red paint (1m×1m) was immersed in the sea water for one month at the Small-Stone Island (Weihai, China) in September, 2019. The paint was full of marine creatures with different sizes of barnacles on it. Sludge and weeds were wiped out with a trowel, followed by washing with sea water, barnacle footprints exposed. Keep washing with sea water to remove dirties on the paint and the surface of footprints. Then sterile distilled water was used to thoroughly wash the footprints and the paint surface. The footprints were then dried in the air for two days. After that, at the center of each footprint (the footprint type was shown in Figure 2B) a chunk of footprint with a diameter about 2mm was picked up into a EP tube with a pair of sterile clean tweezers. Total 15 such samples were collected from 15 the same type of footprints. Genomic DNA extraction kit (SK1203, Sangon) was used to isolate the total DNA from each chunk of footprint. DNA eluted from each chunk of footprint was dissolved into 35ul TE buffer (10mM Tris, PH7.5; 5mM EDTA). The fifteen above DNA samples were equally mixed (Take 5 ul from each 35-ul sample and mixed into a 75ul mixture sample) into one mixture sample, subjected to PCR (polymerase chain reaction) tests using primers for prokaryotes, eukaryotes, archaea and fungus, respectively (Table 1). PCR was performed using self-made nano-PCR kit [7–9]. PCR products were separated in normal agarose gels, photographed, and correct products were cut and purified with Sangon kit (SK8131). Purified DNA was directly ligated with pMD18-T vector (Takara) for TA cloning according to the pMD18-T kit protocol. Ligated DNA was transformed into competent Trans5α cells (Sangon) and selected on agar plates with the final 150ug/mL ampicillin. White colonies were randomly selected for DNA sequencing in Sangon or QingKe (Qingdao). The sequencing results were checked out with BLAST (https://blast.ncbi.nlm.nih.gov/) for species determination suggestions.

**Figure 1.**
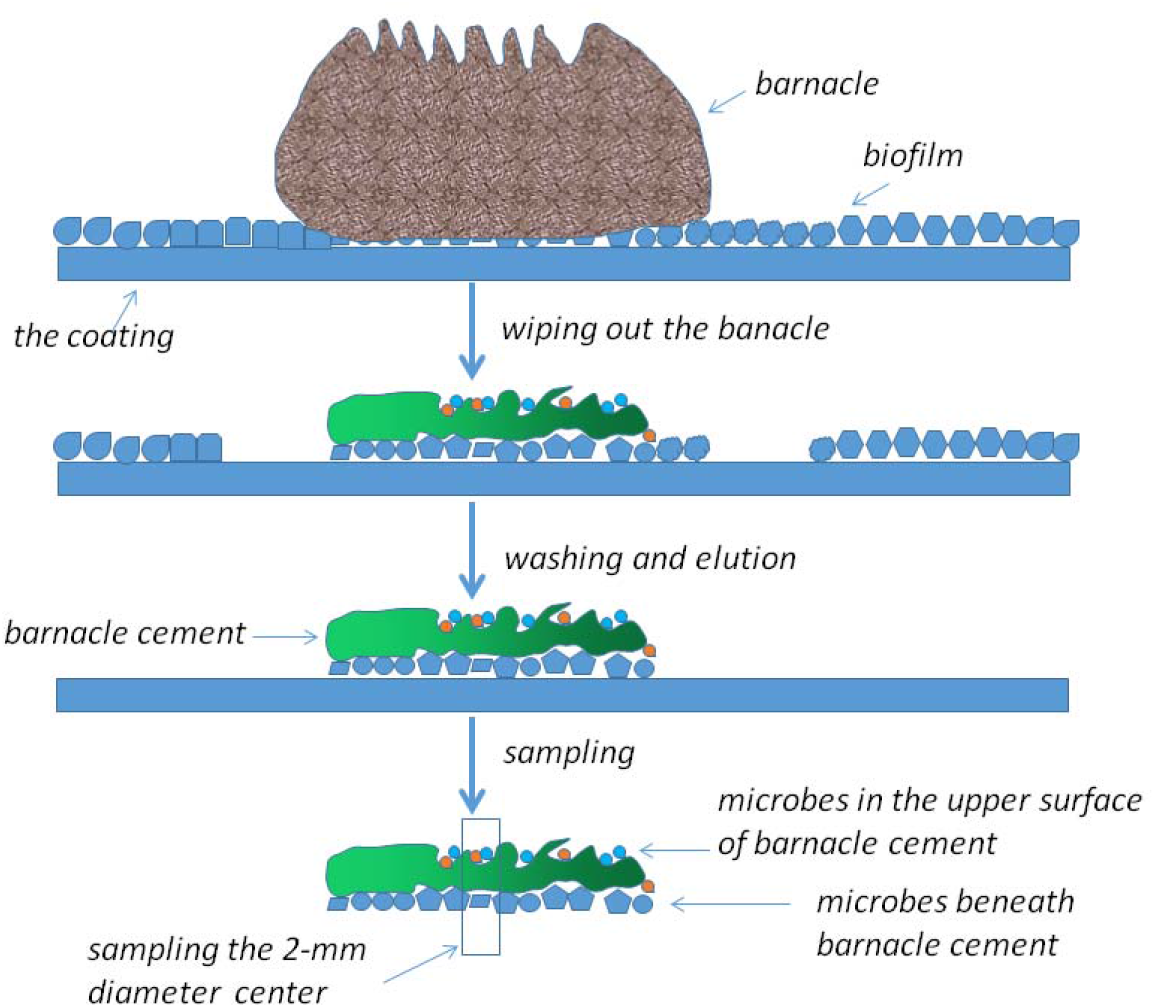
Experimental design of this study

**Figure 2.**
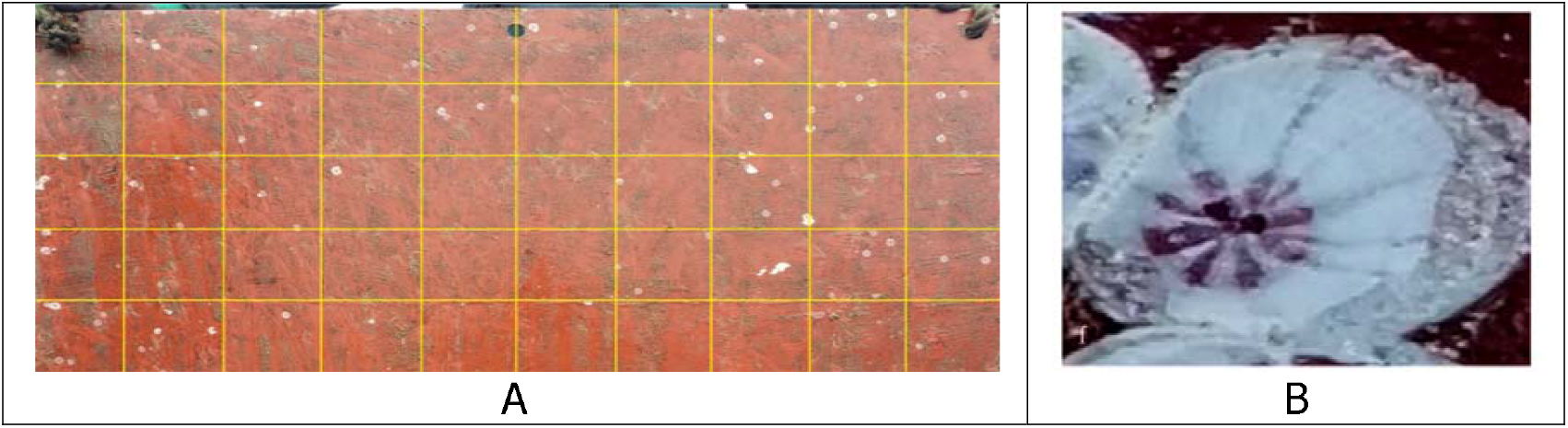
Barnacle footprints on iron-red paints (A) and one example footprint (B). Sampling diameter was about 2 mm in the center of each footprints.

**Table 1.**
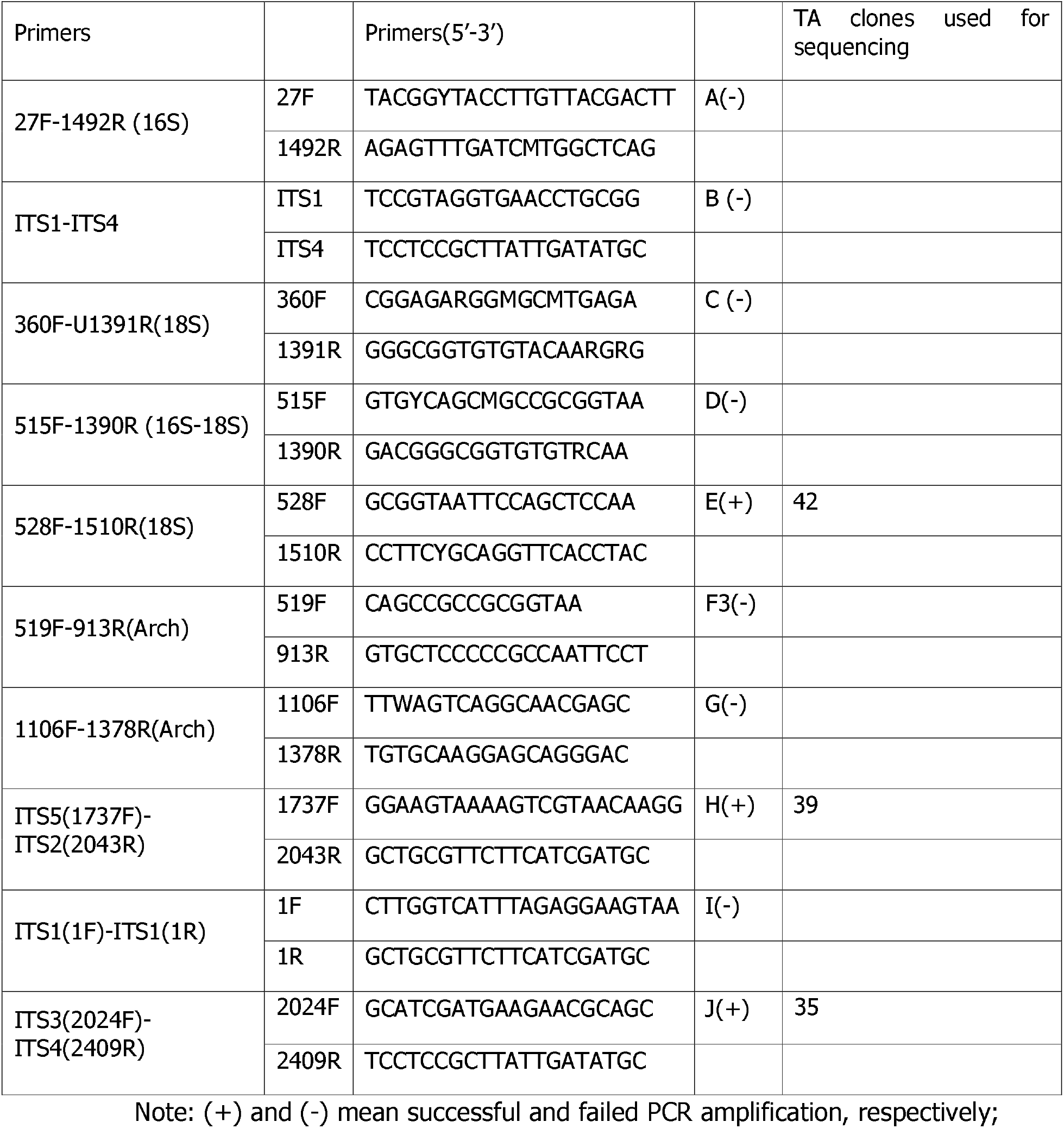
PCR primers

## Results and discussion

For the ten pairs of primers in Table 1, the expected PCR product sizes (bps) are A(1465), B(500-700), C(1032), D(876), E(983), F(395), G(273), H(307), I(200-350), and J(386). In Figure 3, A, B, C, F, and G didn’t get expected sizes of PCR product; D and I failed because their negative controls also showed wrong bands and even repetitions for these experiments still failed. Only E, H and J had decent results and their PCR products were cut and purified for further TA clones and sequencing.

**Figure 3.**
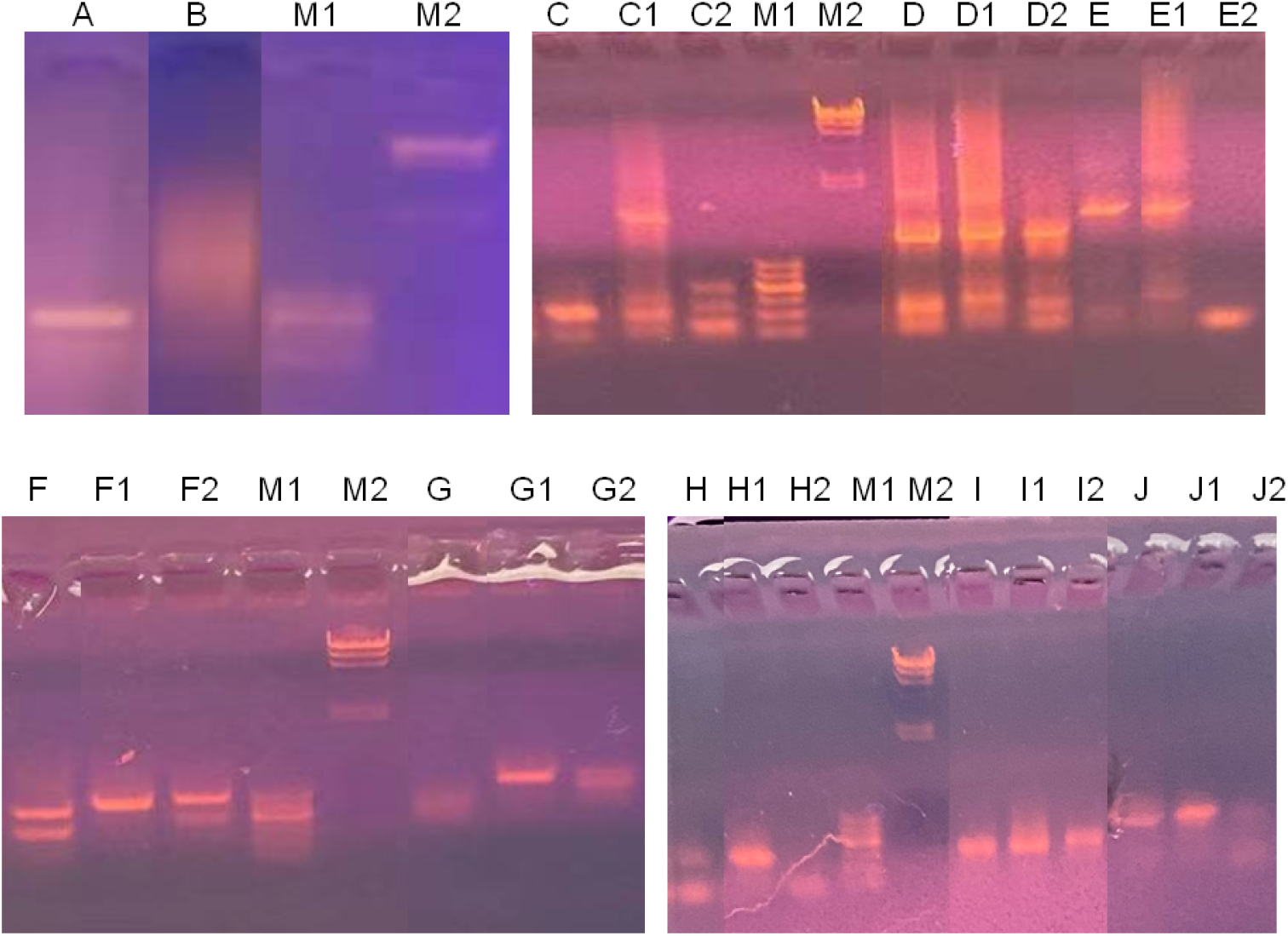
PCR results for A-J samples. M1: markerB with 100, 200, 300, 400, 500 and 600bp bands (Sangon); M2: Lambda/HindIII marker (Sangon); C1, D1, E1, F1, G1, H1, I1 and J1: positive controls for C-J samples, respectively. C2, D2, E2, F2, G2, H2, I2 and J2: negative controls for C-J samples, respectively. All positive controls are the same PCR template made of crude marine biofilm sample; All the negative controls are the same distilled water.

E, H and J TA clones were subjected to Sanger sequencing with 42, 39 and 35 clones, respectively. BLAST (https://blast.ncbi.nlm.nih.gov/) analysis gave suggested species determination. The most frequent species in E, H and J were listed in Table 2. It is interesting that only E and J have a overlapping genus *Aspergillus*, which is known for biofilm formation propensity [10]. Little study has linked these strains in Table 2 with barnacle larva attachment. By this time, microbes beneath the barnacle cement and in the upper surface of it were not separated with each other; future studies shall employ more dedicate approach to specifically pick up microbes beneath the cement.

**Table 2.**
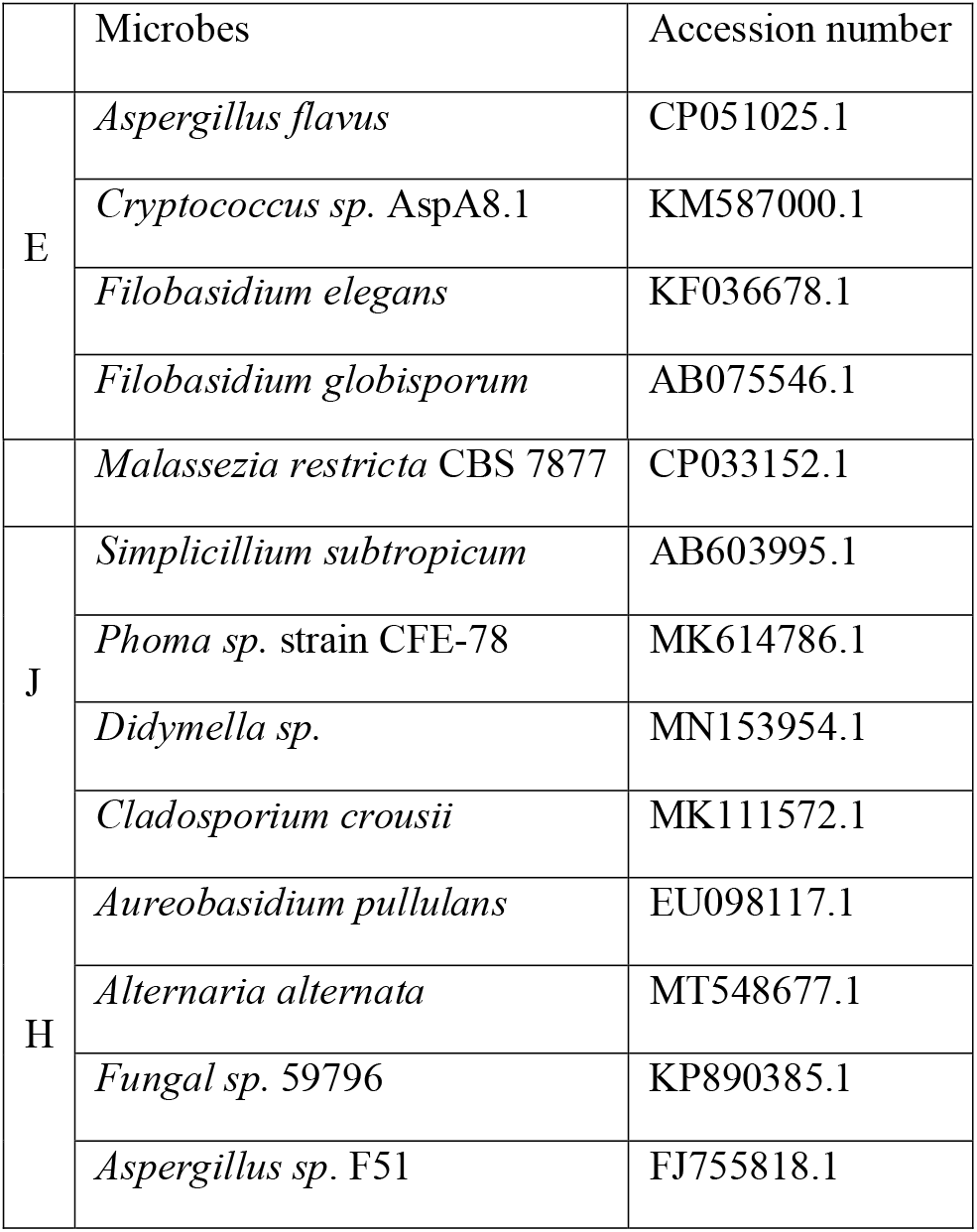
BLAST results for TA clones

This study was not able to locate some common species among E, H and J three groups of data, suggesting that the potential targeted microbe(s) have not been pinpointed yet. Do these different strains have a common feature that can attract barnacle larva? We have no idea about it yet. On the other hand, Figure 1 already pointed out that bacterial contaminants are easy to bring into the experiment, while the nano-PCR kit has very high sensitivity [7–9] to amplify any microbe in the samples in case conservative 16S, 18S and ITS sequences are PCR targets. Once the contaminated microbes have similar amounts as targeted microbes beneath the cement, the whole amplification output would be distorted and the real target microbe information may be lost. Next step we need to precisely pick up the target samples. Besides, it seems that Weihai sea-area has about seven different types of barnacle footprints (data not shown), and delicate experiments would be designed and performed in the near future.

### Implication in student research training

This investigation provided a case study for undergraduate or graduate student to go through a relatively complete laboratory research process: planning, design, sampling, data collection, training of experimental techniques, data statistics, and manuscript/report writing.

## Conclusions

This study provided a novel angle to tackle the question whether barnacle larva attach on a biofilm area with specific microbes. From the coatings immersed in th sea water for some time, barnacles on the coating surface can be obtained, followed by barnacle footprints. It is postulated that beneath the barnacle footprint, the wanted specific microbes may be hidden. This study gave a preliminary test and a name list of microbes (mainly fungi) was obtained. However, whether they are really wanted target microbes largely depends on whether they possess some common features that can attract barnacle cyprids.

## Acknowledgments

2018 Weihai Scientific Innovation Project (grant number 2018HW13); the Key research and development plan of Shandong Province (grant number 2016GSF115022); the Natural Science Foundation of Shandong Province (grant number ZR2018MC002); The author thanks Pengju Gao, Hongwei Shi and Qianqian Liu for their help in sampling.

